# Honey bees perform fine-scale detailing that continuously reduces comb area after nest expansion

**DOI:** 10.1101/2023.09.11.557237

**Authors:** Claire S. Bailey, Peter R. Marting, Michael L. Smith

## Abstract

Honey bees are renowned architects. The workers use expensive wax secretions to build their nests, which reach a mature, seemingly steady state, relatively quickly. After nest expansion is complete, workers do not tear down combs completely and begin anew, but there is the possibility they may make subtle changes like adding, removing, and repositioning existing wax. Previous work has focused on nest initiation and nest expansion, but here we focus on mature nests that have reached a steady-state. To investigate subtle changes to comb shape over time, we tracked six colonies from nest initiation through maturity (211 days), photographing their combs every 1-2 weeks. By aligning comb images over time, we show that workers continuously remove wax from the comb edges, thereby reducing total nest area over time. All six colonies trimmed comb edges, and 98.3% of combs were reduced (n = 59). Comb reduction began once workers stopped expanding their nests and continued throughout the experiment. The extent to which a comb was reduced did not correlate with its position within the nest, comb perimeter, or comb area. It is possible that workers use this removed wax as a reserve wax source, though this remains untested. These results show that the superorganism nest is not static; workers are constantly interacting with their nest, and altering it, even after nest expansion is complete.

## Introduction

The honey bee nest is an essential multi-purpose organ of the superorganism – the hexagonal cells are used for stockpiling food, rearing brood, and form a substrate upon which workers interact (Wheeler 1928; Seeley 1989; Hölldobler and Wilson 2009; Smith 2021). When a swarm of bees moves into an empty cavity, the workers immediately, and feverishly, begin building their nest of wax comb. The initial growth phase lasts approximately 45 days, during which the colony constructs 75-90% of its first-year comb (Lee and Winston 1985; Smith et al. 2016; Marting et al. 2023). After this explosive period, comb growth slows and eventually stops. Comb growth is critically important for a colony’s initial success, but it is also tightly regulated to prevent wasted resources. Given that wax is expensive for bees to produce (7:1 honey:wax; (Weiss 1965; Hepburn et al. 1984)), workers will only further expand their nest after the initial growth phase if two conditions are met: (1) plentiful incoming nectar, and (2) lack of storage space (Pratt 1999, 2004).

The nest of the Western honey bee, *Apis mellifera*, is made up of multiple parallel combs, and each comb is made of thousands of hexagonal cells (Seeley and Morse 1976; Smith et al. 2021, 2023). Workers start construction with a spine of wax which they mold into cells (Huber 1814; von Frisch 1974; Pratt 2000; Franklin et al. 2022). As workers construct cell walls from the wax base, they maintain a border of unshaped wax along the outer edge of the comb (Franklin et al., 2022). This “leading edge” is visible in combs that are actively being built, and serves as a growth front where workers can deposit wax and shape it into new hexagonal cells (Casteel 1912; Franklin et al. 2022; Gallo et al. 2023).

Once a colony builds its nest, the cells are considered fixed. Combs will darken and stiffen with successive brood generations (Zhang et al. 2010), but workers do not typically tear down combs and reshape cells, with the exception of queen cells, which are temporary structures (Allen 1965). The contents of the nest, however, are inherently dynamic, with cells being used for different contents throughout the colony’s life (Smith et al. 2015, 2016). A nest’s total comb area can also appear to fluctuate when colonies reach maturity (Smith et al. 2016), though this could be a methodological artifact (e.g., hand-tracing comb area onto transparent sheets, which are then scanned and digitized). Here, we use high-resolution photography, which makes it possible to capture subtle changes in comb area throughout the colony’s lifetime.

Surprisingly, little is known about comb maintenance once the nest has reached a mature steady-state. The leading edge is particularly interesting, because it has no known function once comb construction is complete, and workers are known to be frugal with their wax. Why would workers maintain an unmolded wax border in a mature nest that is no longer growing? To examine how combs change throughout a colony’s life, we photographed six colonies living in three-dimensional nests every 1-2 weeks from nest initiation through maturity (59 natural-built combs over 211 days total). Aligning 1280 comb images over time, we observed that the leading edge was retracted once nest expansion was complete, but also that this reduction at the comb edge persisted, continuously, in all colonies and 98.3% of combs. Therefore, the wax combs in the honey bee nest are not as fixed as they initially appear.

## Methods

To observe the process of comb growth and maturation, we installed six *A. mellifera* colonies into 10-frame wooden Langstroth hive boxes (38 × 47 × 25 cm) at Auburn University (32° 40’ 26.62’’ N, 85° 30’ 43.55’’ W). Each nest box contained 10 wooden bee frames (20 × 43 cm), without wire supports or wax foundation, so the colonies were free to build their combs naturally within the plane of each bee frame (hereafter, frames). Frames were oriented perpendicular to the nest entrance (frame 1: furthest from entrance, frame 10: closest to nest entrance), and colonies could initiate their nest on any frame within the nest box. Colonies were initiated as artificial swarms (“packages”), each containing 10,000 workers and a mated queen (Gardner Apiaries, Baxley, GA). We fed each swarm sucrose solution (1:1 sugar:water, by volume) for 72 hours pre-installation to induce worker engorgement, which mimics natural swarm conditions (Combs 1972; Seely 2011). On Sunday 4 April 2021, we installed each swarm into a bee box and tracked their comb growth until 1 November 2021 (211 days). Each colony received a 2L sucrose solution feeder on 5 April 2021, but otherwise were given no supplemental feed throughout the experiment.

### Data Collection

To track comb growth over time, we photographed each frame of each box using a high-resolution camera mounted on a custom rig (camera: Nikon Z50, aperture: f7.1, shutter speed: 1/80s, focal length: 135mm; photography rig: 348 × 66 × 44 cm, with LED lighting, and diffusive fabric covering). To obtain an unobstructed view of the comb, we gently brushed the bees off each frame before photographing the comb, and then immediately returned the frame to the colony. From 12 April to 23 June 2021, we photographed colonies weekly; from 23 June to 1 November 2021, we photographed colonies every two weeks as comb construction slowed and changes in comb shape were subtle.

### Data Analysis

To measure the comb area in each image, we trained a neural network to classify each image pixel as either: comb, wooden frame, or background. We first annotated a subset of images by hand, using the Labelbox platform. These annotations were then used to build our model with DeepLabv3, a ResNet-50 backbone, and a subset of the COCO dataset, implemented in torchvision (Lin et al. 2014; He et al. 2016; TorchVision maintainers and Contributors 2016; Chen et al. 2017). The model contained 99 human-annotated images, and 33 images in the validation set. To train the model, we used 100 epochs of stochastic gradient descent, a learning rate of 0.05, and a momentum value of 0.5. For further details on model parameters, see supplemental information in Marting et al. (2023). To detect changes in comb area over time (pixel to mm conversion: 9.96px = 1mm), we used keypoint detection to align sequential images of each frame. Each image output was verified by a human to confirm accurate labeling and alignment.

We established a mature phase within the nest’s life cycle by determining when workers were no longer expanding their comb. For each frame, we calculated: maximum comb area, date of maximum comb area, distance to nest entrance, nest initiation position, and comb perimeter. To determine whether the comb shape was influenced by its position within the nest architecture, we differentiated between combs in the middle of the nest (center) and the periphery of the nest (edge), independent of where the nest was built within the nest box (i.e., at the center or the edge of the box). We identified the nest initiation position as the frame upon which the most comb was built and assigned each frame a rank based on its comb area (rank 1 having the most comb). We defined the top three frames (rank 1-3) as the nest center, and the nest edge as the frame with the smallest nonzero comb area. For each comb, we calculated its perimeter using the using the arc length contour feature in OpenCV (Bradski, 2000).

After reaching a maximum nest area, only one colony, colony number 6, further expanded its nest during the experiment. Therefore, the growth patterns of colony 6 are out of sync with the other experimental colonies. Given that this was the only colony to exhibit additional nest growth, it was dropped from the subsequent analyses. The other five colonies did not substantially grow after the initial growth phase, and so we focus on these five colonies to show how combs change in a mature nest (i.e., one that is no longer growing).

### Statistical Analyses

Statistical analyses were conducted using Python version 3.10 (Python Software Foundation), and the statsmodels, pandas, and numpy packages (Seabold and Perktold 2010; McKinney 2011; Harris et al. 2020). We used an unpaired t-test to determine differences between comb reduction at the center and edge of colonies. To test if comb reduction differed significantly between frame location, we used an ANOVA test. We used linear regression to determine if there was a relationship between comb reduction and the size of the comb.

## Results

Colonies reached their maximum nest area after 80 ± 18 days (8 June to 13 July 2021) at which point growth stabilized as workers stopped expanding the nest. The maximum nest area was 1775.9 ± 97.9 cm^2^ and the maximum comb area per frame was 1778.9 ± 255.3 mm^2^ (note: nest area is reported in cm^2^; comb area in mm^2^). At the nest-level, we define the “growing” phase as when nest area is increasing, and the “mature” phase as when the nest area reaches a plateau. Within a nest, however, not all combs reach their maximum area on the same day, so a nest can contain growing combs and mature combs simultaneously. Nevertheless, the combs which have not yet reached their maximum area (i.e., are still growing) are almost complete (>90% of their maximum area). Therefore, the nest-level estimate of a “mature” phase is reliable even though there is some variation at the comb-level.

Once nests reached the mature phase (i.e., were no longer expanding), nest area and comb area decreased (Fig. 1a, b). To confirm that this was not a methodological artifact, we aligned sequential images of the same comb, and found that workers make subtle alterations that reduced the leading edge at the tip of the comb (Fig. 1c). This same pattern of comb reduction also occurred in the colony that was excluded from our analyses (colony 6 had a secondary growth phase); whenever this colony was not growing, the combs showed a reduction in area that matched our observations from the other five colonies (see below).

**Fig. 1.**
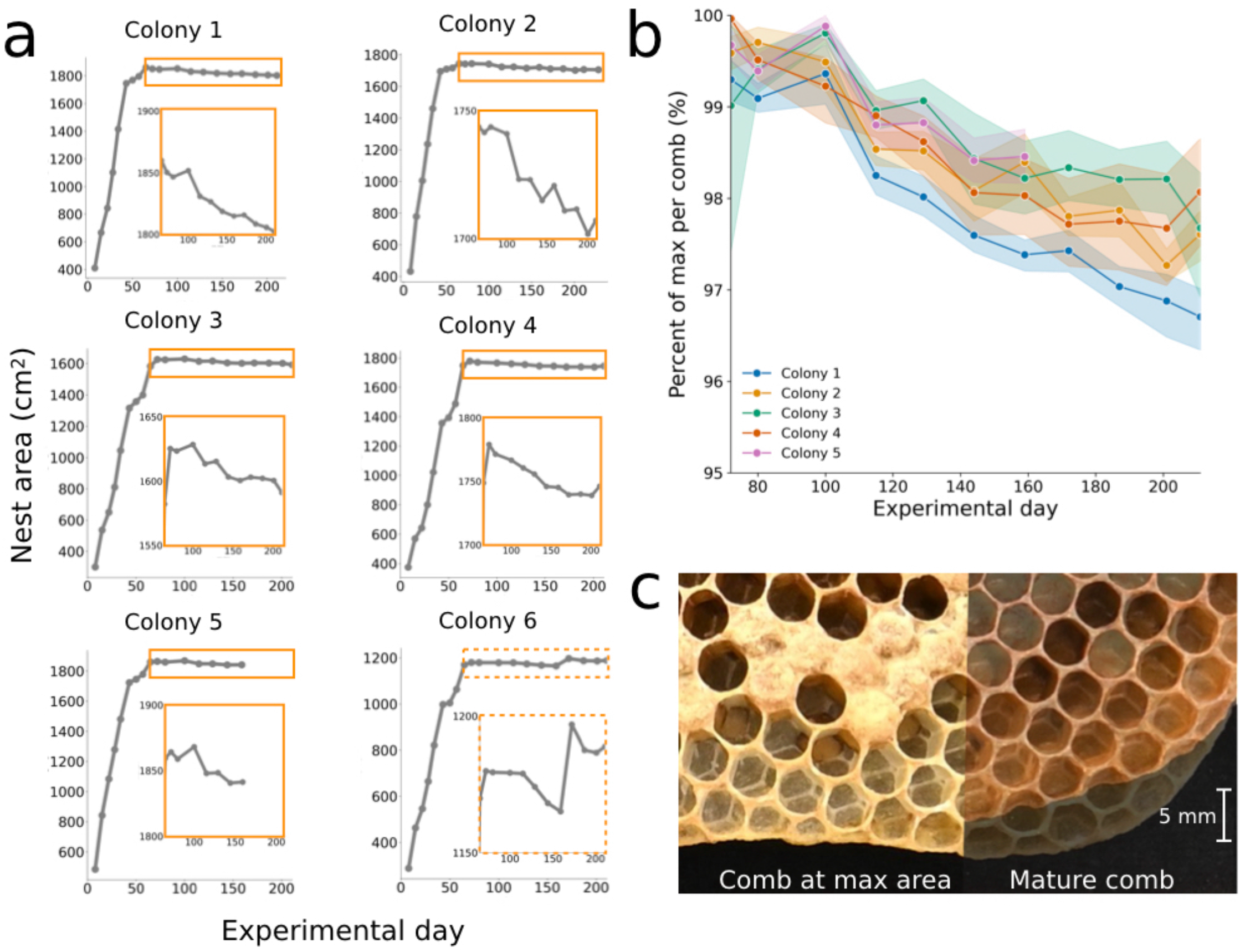
Honey bees reduce their comb area over time. (a) Total nest area from all six colonies throughout the experiment, including the initial growth phase, maximum nest area, and maturity. Orange insert highlights nest area reduction during the mature phase (dotted orange line for colony 6, which shows a secondary growth phase). (b) The percent of maximum comb area during the mature phase. Color denotes colony ID. (c) Overlaid comb images to compare comb at max area (experimental day 65) versus after comb has been reduced (experimental day 211)

Comb reduction was pervasive; it occurred in all six colonies, and 98.3% of combs (n = 59 combs; 58 with reduced area). All data reported hereafter excludes colony 6. Nest area was reduced by 38.3 ± 11.6 cm^2^, accounting for 2.15% ± 0.62% of total nest area. Individual combs were reduced by 41.3 ± 11.0 mm^2^, comprising 2.30% ± 0.59% of individual comb area. At the comb level, workers reduced comb for the first time on day 50 (24 May 2021). At the nest level, comb reduction was first detected on day 72 (15 June 2021), and continued, continuously, until the end of the experiment on day 211 (1 November 2021). Therefore, comb reduction was not a one-time removal of excess wax from the leading edge, or a simple reshaping of the comb tip, but rather a continuous process which reduced nest area at a rate of 0.33 ± 0.09 cm^2^ per day (Fig. 1).

Next, we explored the relationship between comb reduction and nest structure, including: the comb’s position in the nest box (frame number), relative nest location (center versus edge of the nest), comb perimeter, and comb area (Fig. 2). We found no significant difference in comb reduction with respect to: position in the nest box (frame number; ANOVA, p = 0.49, Fig. 2a), position in the nest (nest center versus edge; T-test, p = 0.23, Fig. 2b), comb perimeter (Linear regression, p = 0.12, Fig. 2c), or comb area (Linear regression, p = 0.23, Fig. 2d). Therefore, workers reduce comb independently of the comb’s location in the nest box, distance from the nest entrance, position in the nest, shape of the comb, or size of the comb.

**Fig. 2.**
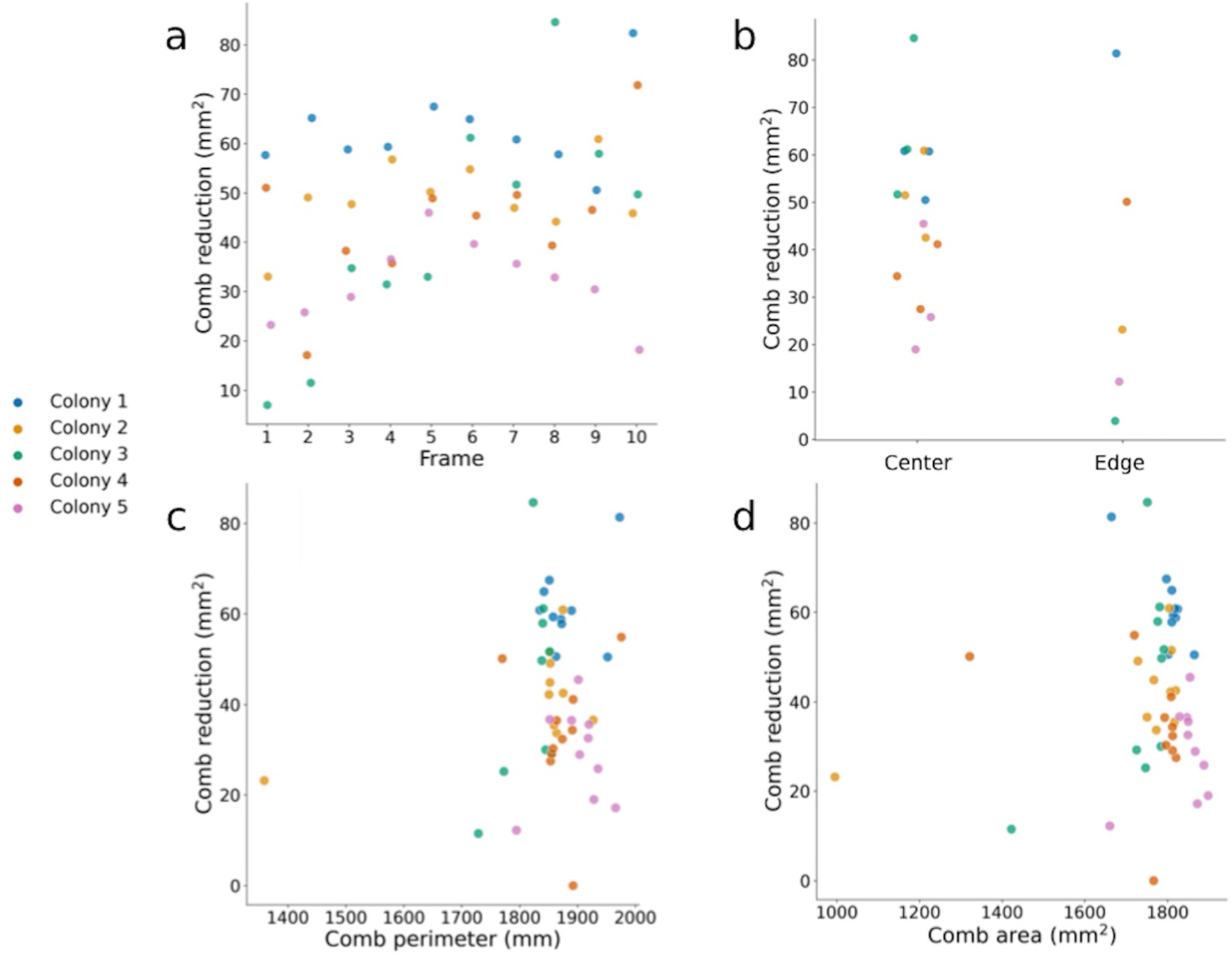
Comb reduction is unrelated to multiple metrics of nest structure. (a) Comb reduction is unrelated to its position in the nest box (frame 1: furthest from nest entrance, frame 10: closest to nest entrance). (b) Comb reduction was no different for combs located at the center versus edge of the nest (the three largest combs are defined as the nest center; the smallest nonzero comb is defined as the edge). (c, d) Comb reduction is unrelated to the comb’s perimeter (c) or area (d). Color denotes colony ID

We also compared morphological features of growing combs and mature combs (Fig. 3). The leading edge of a growing comb was more prominent than in mature comb; as comb matures, the leading edge was removed, and the edge of the comb was smoothed (Fig. 3a). Viewing mature comb ventrally (Fig. 3b), the bottom of the cells were nearly exposed, and one can view the midrib that divides the two sides of the comb. In the cross section, morphological differences were especially clear (Fig. 3c); growing comb formed a sharp point of unfinished and partially-finished cells at the tip, versus mature comb which had a rounded edge of full-depth cells.

**Fig. 3.**
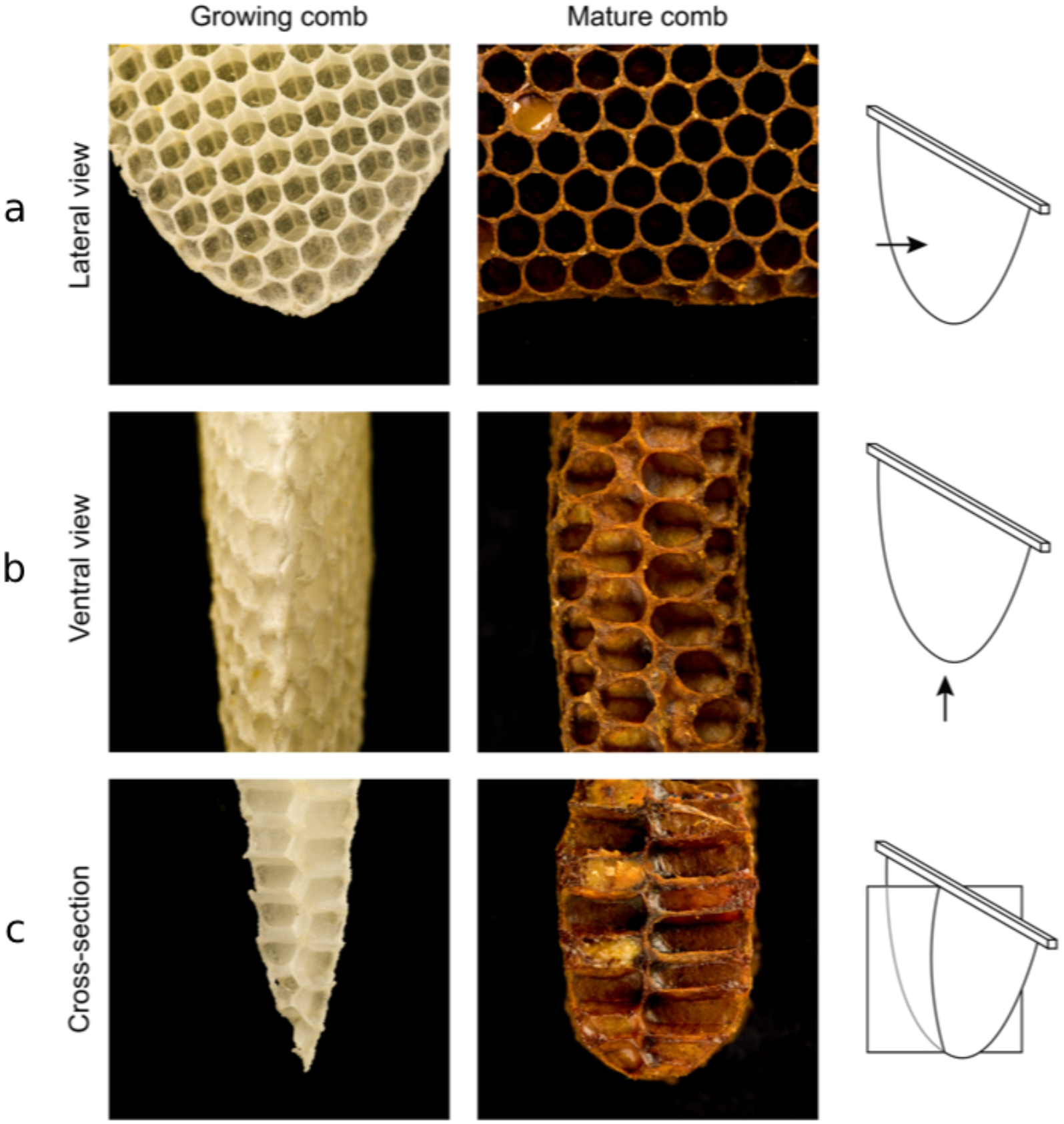
Pieces of growing and mature comb from three different perspectives: (a) lateral, (b) ventral, and (c) cross-section. Illustration at right shows comb built upon a wooden frame, with arrow (or intersecting plane) denoting viewpoint. The mature comb images shown here were collected at the end of the experiment

Finally, we examined colony 6, to assess comb reduction in a colony with secondary growth (Fig. 4). Colony 6 reaches its initial mature phase at the same time as the other five colonies, and similarly experiences comb reduction (Fig. 1a, 4a-c). However, secondary growth is observed on experimental day 172, with a sudden increase in comb area (orange insert in Fig. 4a, 4d). Once reaching its maximum, however, comb area reduces again (Fig. 4a, 4e). Therefore, comb reduction is ubiquitous in the nest, resuming after bouts of comb expansion. The leading edge was removed after the initial growth phase, as seen in the other colonies. During the secondary growth, the leading edge was added back to the comb (Fig. 4d). This leading edge is again removed during the secondary comb reduction (Fig.4, b-e).

**Fig. 4.**
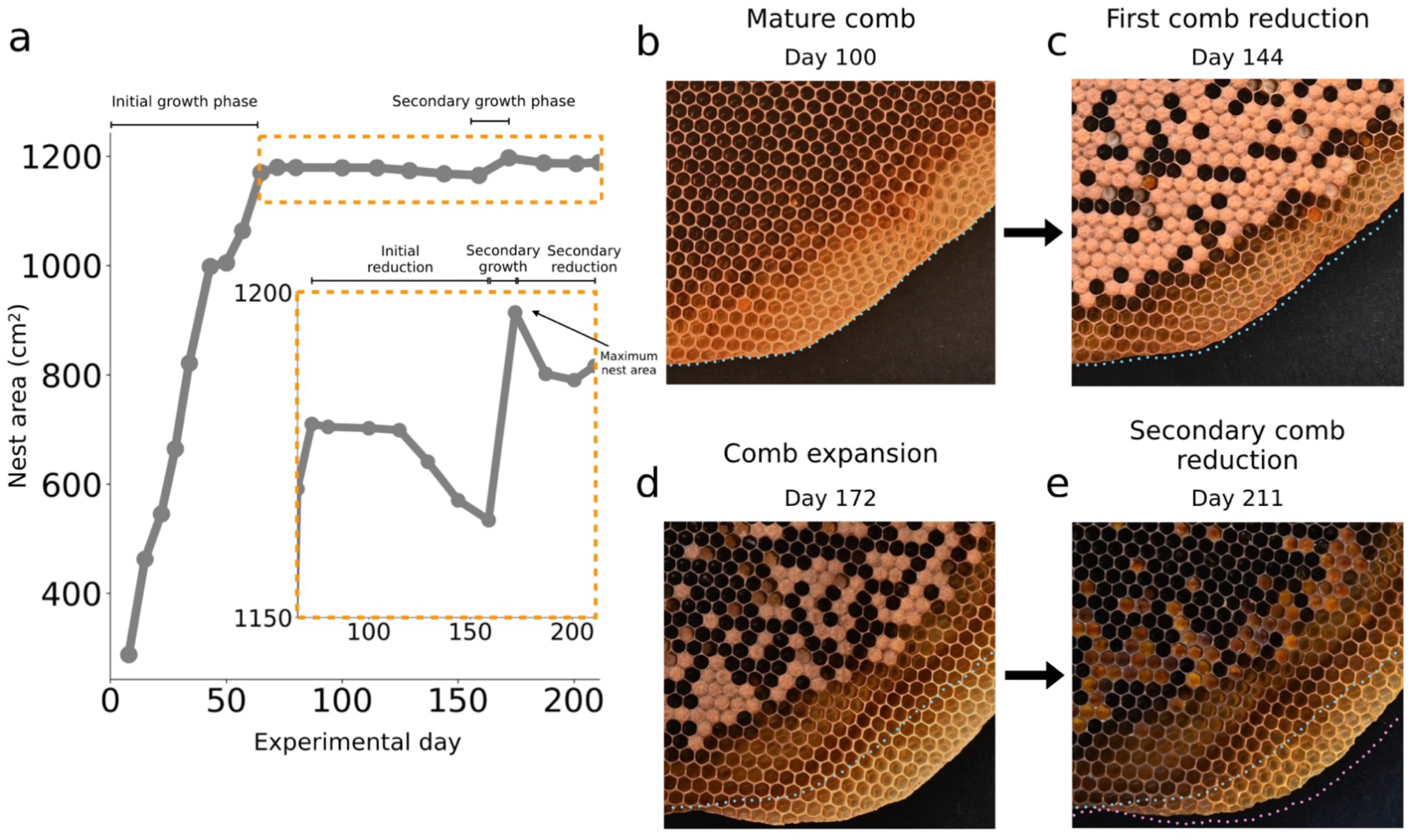
Comb reduction also occurs in colony 6, which had a secondary growth phase. (a) Total nest area for colony 6, with initial and secondary growth/reduction phases labelled. Orange insert highlights nest reduction during the same time period as Fig. 1a. (b-e) Aligned comb images over time highlight phases of comb reduction and comb expansion. Blue dotted line shows comb outline on day 100 ; pink dotted line shows comb outline on day 172 (maximum comb area)

## Discussion

We show that honey bee workers continuously alter their combs, far beyond the initial growth phase. Across all colonies and the majority of combs, we observed a slight, but continuous, reduction in comb area that spanned multiple months (8 June – 1 November 2021). Aligning comb images over time, we find that the comb is reduced at the edges (Fig. 1). The thin and delicate leading edge flattens out and loses its point, thereby forming individual cells rather than a collection of excess wax (Fig. 3). This pattern is observed even in colonies experiencing multiple growth phases (Fig. 4). While the total reduced nest area was minimal (38.3 ± 11.6 cm^2^; 2.15 ± 0.62 %; 0.33 ± 0.09 cm^2^ per day) some combs were reduced by up to 4%. Why certain combs were reduced more than other combs remains unknown, but we did not find a correlation with distance from the nest entrance, location within the nest, comb perimeter, or comb area (Fig. 2). These findings demonstrate the fine-scale dynamic nature of comb construction and nest maintenance, even in mature nests that may initially seem to be stable.

Presumably, bees chew this comb away, given that workers use their mandibles to manipulate wax (Casteel 1912; Siefert et al. 2021). If workers were simply trimming away the leading edge, we would predict: (1) comb reduction to stop once workers had eliminated the excess wax that was not incorporated into a complete cell, and (2) a positive relationship between perimeter and comb reduction. However, neither of these predictions were met. We found that comb reduction was continuous and did not correlate with a comb’s perimeter (Fig.1, 2c). Therefore, comb reduction is more than just removing wax that is left over from the initial growth phase.

Given that honey bees are frugal with their precious resources, it is possible that workers use the tip of the comb as a reserve wax source (e.g., for repairs, capping brood, bolstering existing walls, and capping honey). This could reduce the need for workers to activate their own wax glands – a potential honey-saving technique. Workers of the dwarf honey bee, *A. florea*, for example, are known to steal wax from abandoned colonies (Hepburn, 2010), though this behavior has not been reported in *A. mellifera*. It is possible that workers may be repurposing, or recycling, wax resources within their own colony. To confirm this, one would need to observe comb reduction directly, perhaps using long-term recordings (e.g., Siefert et al. 2021). If bees are recycling wax within their nest, this would be a newly discovered behavior in an already-impressive animal architect.

Of the six colonies, only one had a secondary period of comb expansion, which was observed on experimental day 172 (23 September 2021), 14 weeks after reaching its initial maximum nest area. This secondary growth was not for building reproductive nest architecture, but likely occurred due to plentiful nectar and limited storage space, which is known to induce comb growth (Pratt 1999, 2004). Colony 6 was excluded from our analyses because it was out of sync with the other five colonies. Data from this colony, however, can be used to confirm the general trends we observed. This colony also exhibited comb reduction when the nest was not growing (Fig. 4). Furthermore, workers manipulate the leading edge before and after nest expansion (Fig. 4, b-e). Therefore, the leading edge is likely important not just for the initial growth phase (Franklin et al. 2022), but also when colonies undergo additional nest expansion. Whether and how the leading edge may signal comb expansion to workers is unknown, but it is an intriguing area for future work.

Honey bees are a well-studied system, so it is surprising that comb reduction has not, to our knowledge, been previously reported. This is likely for two reasons. First, to detect these subtle changes, one needs repeated high-resolution images of natural-built combs, and to align them. Previous work has typically used transparency sheets, or estimates using grid-squares (Pratt 1999; Rangel and Seeley 2012; Smith et al. 2016), which are relatively low-resolution estimates, and therefore might not detect small changes in comb area. Second, comb reduction was only observed once the nest was no longer expanding. Previous work has focused on the process of nest expansion, either in the initial stage of moving into a new nest cavity (Lee and Winston 1985; Rangel and Seeley 2012; Smith et al. 2016; Marting et al. 2023), or when a colony must decide whether to invest in additional storage space (Pratt 1998, 1999, 2004), not on the combs in their mature steady-state. Our work shows that, given the correct tools, we can analyze the entire life cycle of a nest, to better understand the dynamic nature of this essential structure. Comb reduction shows that workers are always interacting with, and altering, their nests. Like cells in the human body, constantly dying and regenerating, the superorganism nest is dynamic.

## Acknowledgements

We thank Prathibha Chandran, Ben Koger, Ethan Rowe, Michelle Simpson, and Maritza Spott for help with data collection and processing. Clair Dickinson, Erica Maul, and Olivia Smith provided helpful feedback on the manuscript. We also thank Davis Arthur, Eileen Bailey, and David Bailey for their proofreading and support.

